# A Rapid and Modular Nanobody Assay for Plug-and-Play Antigen Detection

**DOI:** 10.1101/2025.03.01.640988

**Authors:** N. Rebecca Kang, John R. Biondo, Caitlin E. Sharpes, Katherine A. Rhea, Padric M. Garden, Juan J. Jaramillo Montezco, Alina Ringaci, Mark W. Grinstaff, Daniel A. Phillips, Aleksandr E. Miklos, Alexander A. Green

**Affiliations:** Department of Biomedical Engineering, Boston University, Boston, MA, USA; Biological Design Center, Boston University, Boston, MA, USA; US Army DEVCOM Chemical Biological Center, Edgewood, MD, USA; Precise Systems, Lexington Park, MD, USA; Department of Chemistry, Boston University, Boston, MA, USA; Department of Material Science and Engineering, Boston University, Boston, MA, USA; Molecular Biology, Cell Biology & Biochemistry Program, Graduate School of Arts and Sciences, Boston University, Boston, MA, USA

**Keywords:** Antigen Detection, Nanobodies, Coiled Coils, Lateral Flow Assays, Cell-Free Protein Synthesis, Modular Design

## Abstract

Rapid and portable antigen detection is essential for managing infectious diseases and responding to toxic exposures, yet current methods face significant limitations. Highly sensitive platforms like the Enzyme-Linked Immunosorbent Assay (ELISA) are time- and cost-prohibitive for point-of-need detection, while portable options like lateral flow assays (LFAs) require systemic overhauls for new targets. Furthermore, the complex infrastructure, high production costs, and extended timelines for assay development constrain manufacturing of traditional diagnostic platforms in low-resource settings. To address these challenges, we describe the Rapid and Modular Nanobody Assay (RAMONA) as a versatile antigen detection platform that leverages nanobody-coiled coil fusion proteins for modular integration with downstream readout methods. RAMONA merges the portability of LFAs with the benefits of nanobodies, such as their smaller size, improved solubility, and compatibility with cell-free protein synthesis systems, enabling on-demand biomanufacturing and rapid adaptation for diverse targets. We demonstrate assay generalizability through the detection of three distinct protein targets, robustness across various temperatures and incubation periods, and compatibility with saliva samples and cell-free synthesis. Detection occurs in under 30 minutes, with results strongly and positively correlating to ELISA data while requiring minimal resources. Moreover, RAMONA supports multiplexed detection of three antigens simultaneously using orthogonal capture probes. By overcoming several limitations of traditional immunoassays, RAMONA represents a significant advancement in rapid, adaptable, and field-deployable antigen detection technologies.

## Introduction

Early detection is key for controlling the spread of infectious diseases and for effective treatment outcomes following toxic exposure. It is essential that the detection of effector proteins or their precursors is fast and accurate for decisive response measures to be initiated. However, the most sensitive detection assays are often time- and cost-prohibitive, limiting accessibility. For example, the Enzyme-Linked Immunosorbent Assay (ELISA) is an industry benchmark for protein detection diagnostics, but the assay requires trained personnel to perform lengthy incubations and meticulous washing steps.^1^ Alternatively, methods like paper-based lateral flow assays (LFAs) are rapid and portable, yet development of such assays for an emergent threat or disease is not trivial.^2^ Accordingly, there is a pressing need for fast and generalizable antigen detection platforms that can be both manufactured and administered rapidly in low-resource settings in response to new threats.

Antibodies are revolutionizing modern medicine with their ability to specifically bind protein targets for diagnostics and therapeutic applications. Notable technologies like the RAPPID sensor platform are fast, portable and sensitive.^3^–^5^ Yet the inherent bulkiness and poor solubility of antibodies complicate synthesis, making development of new immunoassays more challenging.^6,7^ On the other hand, smaller antibody fragments provide an alternative approach to immunoassay development. Nanobodies are single-domain camelid heavy chain antibodies that are a tenth of the size of conventional antibodies, improving accessibility to epitopes on antigen targets and increasing the ease of genetic manipulation.^8^ Moreover, nanobodies consist of more hydrophilic residues, enhancing solubility and stability.^8^ Consequently, nanobodies show promise for use in on-demand, decentralized biomanufacturing of diagnostic assays.

Cell-free protein synthesis systems are a promising tool to streamline protein production. These membrane-less cocktails of core cell machinery are tailored for protein expression and circumvent many challenges associated with *in vivo* production of proteins such as toxicity.^9^ They can be easily reprogrammed by changing DNA inputs to produce a wide array of proteins, a capability that has been leveraged for high-throughput protein production^10^, on-demand biomolecular manufacturing^11,12^, and portable diagnostic systems^13,14^. Stockpiled cell-free reaction systems have the potential for rapid manufacture of immunoassay components for mass distribution in the face of an emergent epidemic. For instance, DNA encoding newly developed antigen affinity capture components can be easily amplified, distributed, and added to stockpiled “blank” cell-free mixes to produce diagnostic devices against novel biological threats at the point of need.

However, such rapidly deployable diagnostic systems will require methods to interface antigen-sensing components produced by cell-free synthesis reactions with convenient assay readout formats. Site-specific bioconjugation techniques provide an extensive toolkit that could be used to this end. There are three broad categories of site-specific bioconjugation techniques: chemical conjugation, enzymatic conjugation, and self-assembly.^15^ The ideal bioconjugation technique for a field-deployable device should be compatible with cell-free production for integrated component production. Chemical conjugations mainly employ reactive groups like azides and cycloalkynes and require synthetic chemistry for synthesis, which renders them inconvenient to manufacture via translation.^15^ Enzymatic conjugations rely on exogenous enzymes, which could complicate synthesis with concerns such as site selectivity^15^ and the functionality of enzymes in cell-free lysates^16^. Non-covalent self-assembly mechanisms utilize multimeric peptides for conjugation, circumventing synthetic chemistries and exogenous enzymes and thus providing an attractive route for site-specific bioconjugation of cell-free products.

A particularly promising system for this purpose is supramolecular coiled coils. Governed by hydrophobic and electrostatic interactions between inter- and intramolecular side chains, coiled coils form stable and predictable multimeric alpha helical bundles via thoughtfully selected amino acid residues arranged in repeated heptad structures.^17^ Importantly, coiled coils are miniscule peptides that can be easily expressed as fusion proteins with sensing elements such as antibodies and nanobodies with minimal steric hindrance. In support of coiled coils as a method for interchangeable nanobody functionalization, a recent manuscript describes the modular conjugation of nanobodies with fluorescent proteins for microscopy.^18^ Prior to this work, the cell-free expression of nanobody-coiled coil fusion proteins capable of self-assembly has not been demonstrated.

Herein, we report a nanobody-based platform, termed Rapid and Modular Nanobody Assay (RAMONA), that enables fast and generalizable antigen detection (Figure 1). Chimeric nanobody-coiled coil fusion proteins can be functionalized with cognate coil partners bioconjugated to virtually any other molecule for interface with various downstream readout methods. We chose to integrate RAMONA into an LFA-style format for ease of use and eye-readable outputs. We demonstrate that RAMONA is generalizable through the detection of three distinct protein targets: RiVax, Apolipoprotein A1, and Fibrinogen. The system is highly robust, maintaining functionality across a wide range of temperatures and incubation periods. We exemplify compatibility with cell-free production through a sandwich screening assay that identified co-binding cell-free produced nanobodies for all three antigens. In the presence of orthogonal capture probes, RAMONA provides multiplexed capabilities to detect all three antigen targets simultaneously. RAMONA is compatible with saliva and requires less than 30 minutes for detection, with strong and positive correlations with ELISA readouts for 10 fresh saliva samples. Overall, we demonstrate RAMONA as an antigen detection platform that can be produced and performed rapidly, with robustness against temperature and semi-quantitative readout when interfaced with lateral flow readouts.

**Figure 1.**
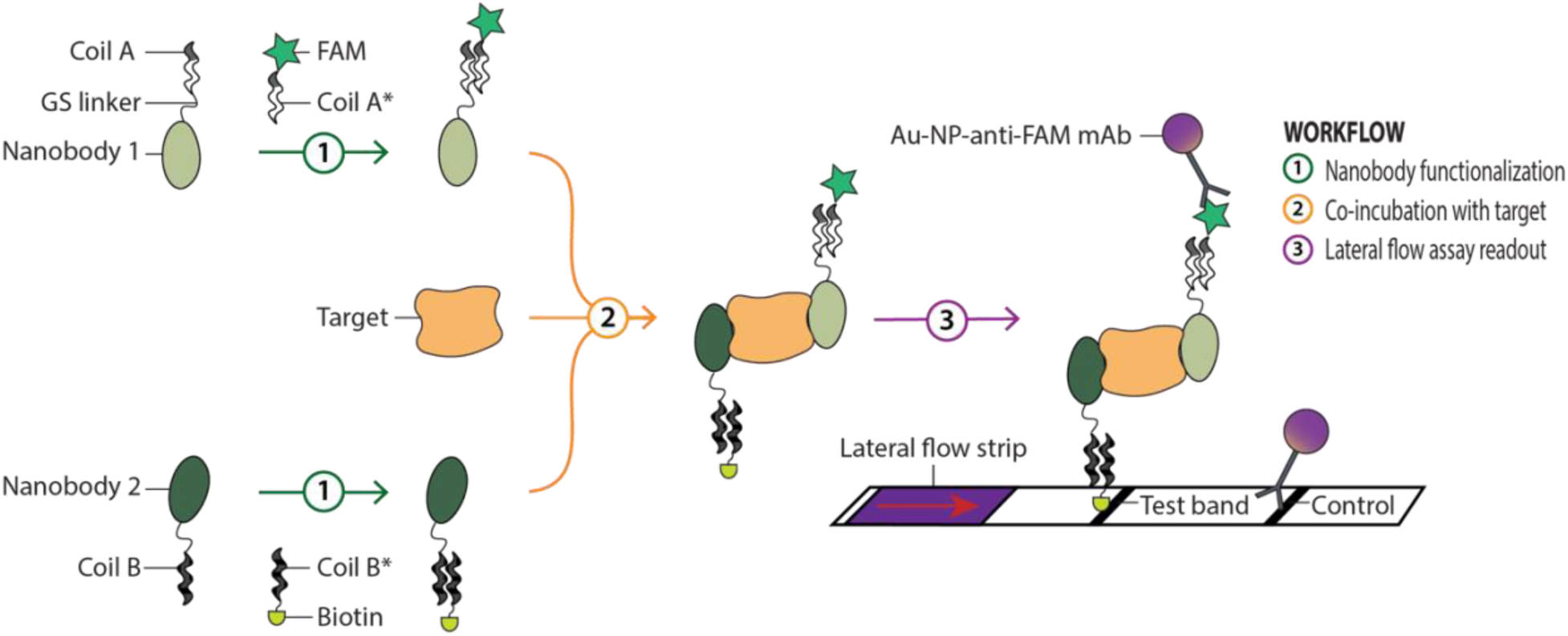
Schematic illustration of Rapid and Modular Nanobody Assay (RAMONA) complex assembly for antigen detection. Nanobodies are produced as fusion proteins tethered to one side of an orthogonal pair of coiled coils via a flexible glycine-serine (GS) linker. They can be functionalized with a reporter (e.g. 6-carboxyfluorescein (FAM)) or capture (e.g. biotin) probe by incubation with cognate coiled coils that were separately bioconjugated to small molecules using copper-free click chemistry. For detection, functionalized nanobody-coiled coil conjugates are co-incubated with the target of interest before interfacing with a downstream output method, demonstrated here in a lateral flow assay format for portability and ease of use.

## Results

We set out to design a compact LFA-based detection system that was both modular and convenient to produce. Lateral flow immunoassays require two main components, analyte recognition elements that bind to the target molecule and capture elements necessary for assay readout. In universal lateral flow strips, the capture elements are often chemical moieties, such as fluorescent dyes and small molecules, that enable the analyte to bind at a specific location along the strip and promote capture of the gold nanoparticles necessary to form a visible band.^19^ Crucially, these capture moieties usually cannot be directly incorporated into the recognition elements during translation, complicating manufacture. In contrast, RAMONA adopts a two-pronged approach to enable flexible and streamlined coupling of recognition elements produced in cell-free systems to capture elements for direct use in LFAs. Firstly, we chose nanobodies instead of antibodies to capitalize on their smaller size^8^, improved solubility^8^, and compatibility with cell-free protein synthesis^20^. Second, we utilized coiled coils, which are only about 4kDa, as a toolkit to functionalize nanobodies for LFA. The specific interactions present within a published collection of coiled coils enables nanobody bioconjugation in an interchangeable, location-specific and ratiometric manner.^21^ RAMONA nanobodies were thus produced as fusion proteins tethered to one side of an orthogonal pair of coiled coils via a flexible glycine-serine linker (Figure 1). Separately, we employed peptide synthesis and click chemistry to generate complementary synthetic coil peptides that contained the necessary LFA capture elements. Nanobody-coil fusions and synthetic coil peptides were then co-incubated to produce functionalized nanobodies. The coiled coils served as a modular interface, effectively decoupling the recognition domains (nanobodies) from the capture domains (chemical moieties). This configuration enables the facile bioconjugation of nanobodies with a wide range of functional molecules, thereby facilitating integration into diverse downstream applications.

Ricin is a highly potent toxin derived from the seeds of the castor bean plant, and its previous use for bioterrorism classifies it as a category B biothreat by the US Centers for Disease Control and Prevention (CDC).^22,23^ Consequently, rapid and early detection methods and treatments for ricin are actively researched, motivating us to assess our novel sandwich immunoassay for ricin detection. Fortunately, a recombinant and catalytically inactive form of ricin known as RiVax is available for safe research and development.^24^ Furthermore, many high-resolution crystal structures are available for the antigen alone and in complex with single nanobodies.^25–28^ We shortlisted six nanobodies (Supplementary Table 1) that exhibited strong binding affinity to RiVax at distinct epitopes for initial testing.^25,26,28^ Figure 2A depicts a representative pair of nanobodies, 202 and 301, predicted to co-bind to RiVax via AlphaFold3 computational modelling. Corroborating this model with reported crystal structures, we reasoned that this would be an ideal pairing for RAMONA characterization as their epitope regions were far enough apart to minimize steric hindrance in the fully assembled RAMONA complex (Figure 2B).

**Figure 2.**
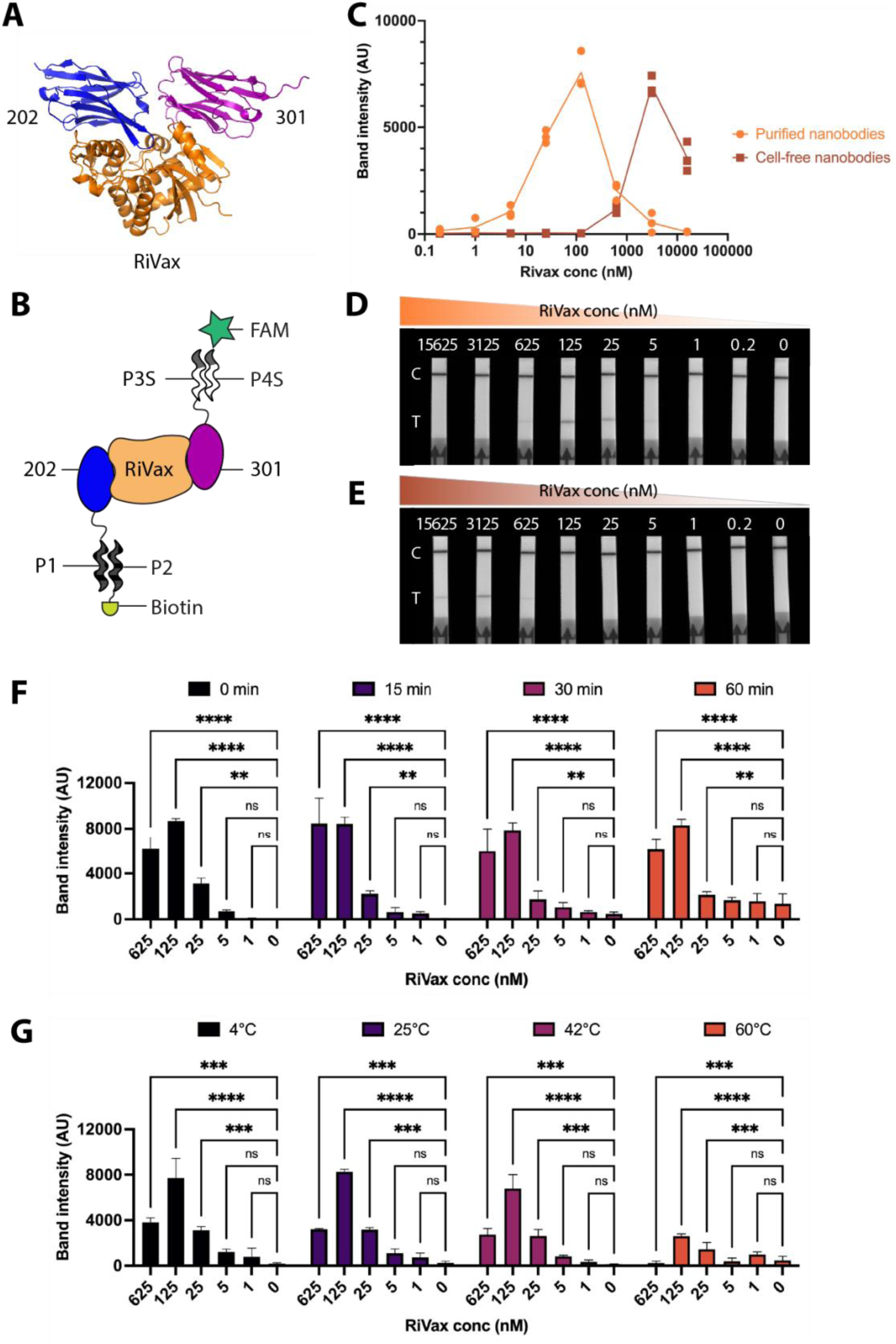
RAMONA complex assembly for RiVax detection. (A) Computational model of nanobodies 202 and 301 sandwich binding with RiVax at separate epitopes. (B) Schematic illustration of sandwich binding of 202 and 301 with Rivax in a fully assembled RAMONA complex. (C) Performance of column chromatography-purified and cell-free produced nanobody-coiled coil conjugates against serially titrated concentrations of RiVax. (D) Representative image of lateral flow strips used to quantify band intensities for the performance of purified nanobody-coiled coil conjugates against titrated RiVax concentrations in (C). “C” indicates control line, and “T” indicates test line. (E) Representative image of lateral flow strips used to quantify band intensities for the performance of nanobody-coiled coil conjugates produced in cell-free systems against titrated RiVax concentrations in (C). “C” indicates control line, and “T” indicates test line. (F) Performance of 100 nM purified RAMONA nanobodies when co-incubated with titrated RiVax concentrations for various time periods before addition to lateral flow strips. (G) Performance of 100nM purified RAMONA nanobodies when co-incubated with titrated RiVax concentrations for 15 minutes at various temperatures before addition to lateral flow strips. Error bars indicate standard deviation across n=3 independent replicate experiments, and statistical significance was determined by two-way ANOVA (adjusted *p* value <0.0001 is denoted by ****, 0.0001 to 0.001 by ***, 0.001 to 0.01 by **, 0.01 to 0.05 by *, and ≥0.05 by ns).

To interface with lateral flow, we synthesized two capture peptides conjugated to biotin or 6-carboxyfluorescein (FAM), which enabled the former to bind to a streptavidin test line in the test strip and the latter to capture anti-FAM antibody-conjugated gold nanoparticles for readout. Histidine-tagged, column chromatography-purified nanobodies were functionalized with these synthetic peptides before being co-incubated with equimolar ratios of RiVax at titrated concentrations or PBS. Lateral flow assays were then run and imaged, and ImageJ was used to quantify the test band intensities on lateral flow strips. A minimum of 62.5 nM of functionalized nanobodies were required for significant LFA distinction between positive and negative results (Supplementary Figure 1). 100 – 200 nM of nanobody-coiled coil conjugates proved optimal for maximizing signal-to-noise ratio while minimizing material usage. Next, we surveyed the ability of 100 nM of purified RAMONA nanobodies to co-bind RiVax over a wide range of concentrations at room temperature (Figure 2C, 2D). Limit of detection was calculated to be 1.28nM.^29^ Band intensity was diminished beyond 125 nM, likely due to excess RiVax causing the Hook effect which hinders nanobody sandwich formation.^30^

Next, we characterized assay performance under various incubation periods and temperature conditions. The co-incubation duration of RiVax with functionalized nanobodies at room temperature did not change the sensitivity of the assay, but increasing incubation times did increase background signal at lower concentrations (Figure 2F). Notably, RAMONA nanobodies co-bound RiVax to produce a signal without any incubation. To probe temperature effects, we co-incubated RiVax with nanobody-coiled coil conjugates for 15 minutes at various temperatures and found that the assay was similarly robust to temperature variations (Figure 2G). Surprisingly, the assay maintained functionality up to 60°C, despite signal intensities being diminished for higher concentrations and increased background noise present.

On-site manufacturing would further support field deployment of a detection platform, so we evaluated if cell-free produced nanobodies were compatible with RAMONA (Figure 3A). We used polymerase chain reaction (PCR)-amplified linear DNA templates of nanobody-coil fusion proteins for overnight cell-free production in Liberum Biotech’s Juice lysate. The next morning, we directly functionalized nanobody-coil fusions within crude lysates with FAM or biotin before repeating the RiVax concentration titration experiment. From Figures 2C and 2E, the limit of detection for cell-free RAMONA nanobodies was calculated as 134 nM.^29^ The Hook effect was again visible past 3125 nM.

**Figure 3.**
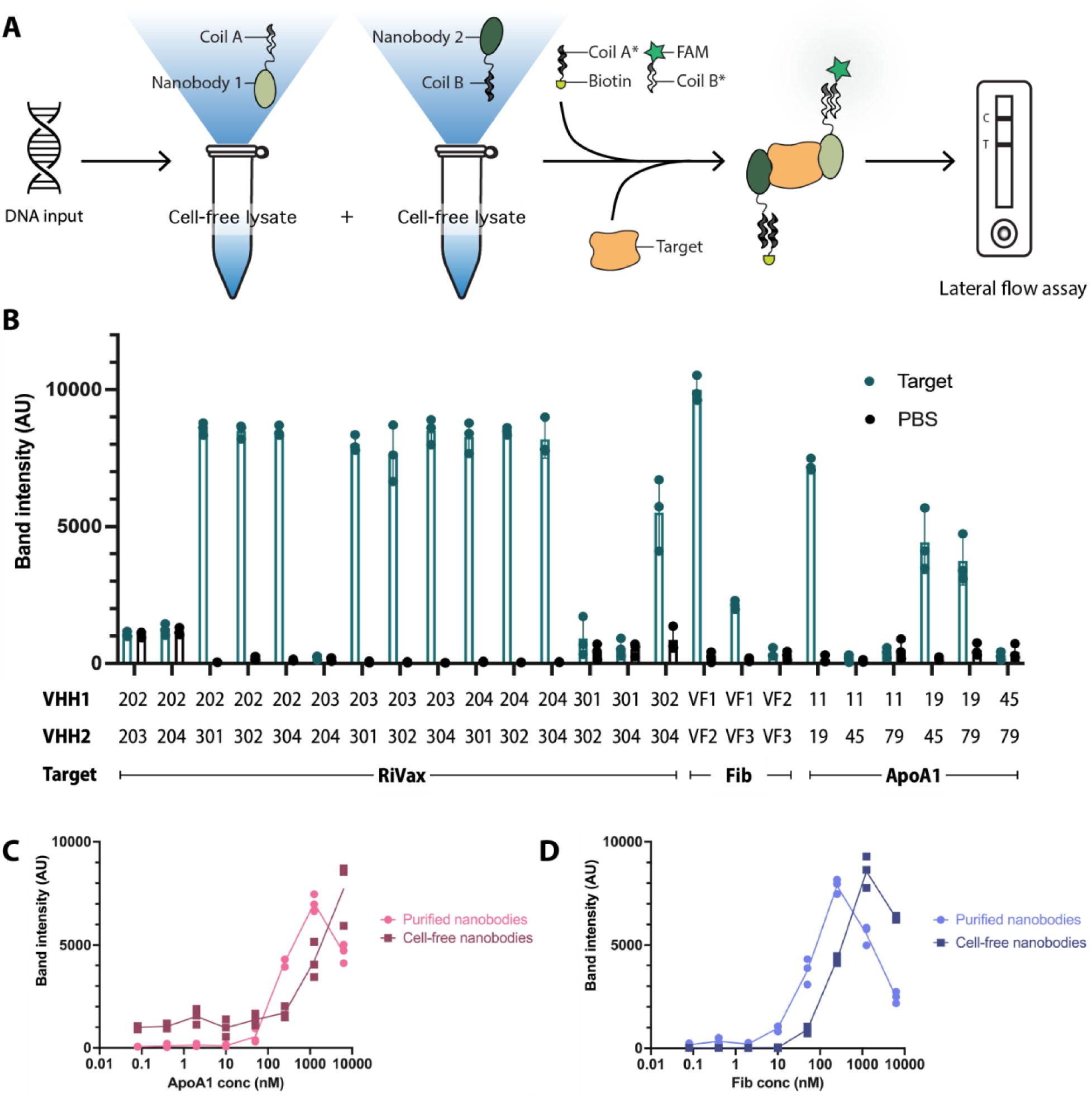
Extending RAMONA capabilities to detect different antigens. (A) Schematic illustration of the workflow for cell-free production of nanobody-coiled coil conjugates for RAMONA. (B) Cell-free screen of nanobody-coiled coil conjugate pairs against various target antigens, prepared as 200nM pure proteins dissolved in PBS. Each antigen was co-incubated with crude cell-free lysates containing the functionalized nanobodies in equivolumetric parts for 15 minutes at room temperature before addition to lateral flow strips for readout. (C) Performance of purified and cell-free produced nanobody-coiled coil conjugates Nb11 and Nb19 against serially titrated concentrations of ApoA1. (D) Performance of purified and cell-free produced nanobody-coiled coil conjugates VF1 and VF2 against serially titrated concentrations of Fib. Error bars indicate standard deviation across n=3 independent replicate experiments, and statistical significance was determined by two-way ANOVA.

Following successful RiVax detection using RAMONA, we validated the platform on additional targets. Recognizing that structural data would not always be available guide nanobody selection, we further postulated that a cell-free-based RAMONA workflow could be employed for rapid functional screens for nanobody co-binders. Apolipoprotein A1 (ApoA1) and Fibrinogen (Fib) were selected as candidate antigens because they are protein biomarkers often implicated in disease^31–35^, with nanobodies targeting them available from the literature, albeit without protein structure models.^36,37^ Using cell-free lysates for nanobody-coil fusion protein production, we tested 15 combinations of 6 RiVax nanobodies, 6 combinations of 4 ApoA1 nanobodies and 3 combinations 3 Fib nanobodies to thoroughly screen for potential co-binders (Figure 3B). As expected, nanobodies 202-204 did not show RiVax-specific co-binding as they shared similar epitope regions. However, they co-bind well with nanobodies 301, 302 and 304, which are known to bind to distinct epitopes on RiVax.^25,26,28^ Surprisingly, 302 and 304 can co-bind, which was previously unclear due to their proximal epitopes. The screen also revealed both expected and novel binding partners for target antigens lacking structural binding data. For ApoA1, nanobodies Nb11 and Nb19 co-bound as predicted, while novel binding partners Nb19 and Nb45, and Nb19 and Nb79 were discovered.^36^ Similarly, nanobodies VF1 and VF2 co-bound Fib as reported in literature, while VF1 and VF3 bound unexpectedly.^37^ The expected nanobody pairs Nb11-Nb19 and VF1-VF2 co-bound targets ApoA1 and Fib best, respectively. We further characterized the ability of these column chromatography-purified versus cell-free produced nanobodies to co-bind their target antigens at titrated concentrations (Figure 3C and 3D). The limits of detection for purified nanobodies and cell-free nanobodies against ApoA1 were calculated as 125nM and 526nM, respectively; the limits of detection for purified nanobodies and cell-free nanobodies against Fib were calculated as 3.13nM and 98.6nM, respectively.^29^

Specificity of nanobody-coiled coil conjugates for their target antigens was next evaluated by co-incubating nanobody sandwich pairs with each of the three targets separately for 15 minutes at room temperature before running lateral flow. All nanobody pairs were highly specific to their respective target antigens, though Fib nanobodies had slightly elevated but statistically insignificant crosstalk signal with the other targets (Supplementary Figure 2). Building upon target specificity, we then investigated if RAMONA can be multiplexed for simultaneous protein detection. We functionalized capture nanobodies for each target with orthogonal probes biotin, 6TAMRA or Cy5, while reporter nanobodies for all targets were functionalized with FAM (Figure 4A). Each target was co-incubated with their nanobody pairs for 15 minutes at room temperature before being mixed in various combinations for lateral flow assays. When a single antigen was exposed to RAMONA nanobodies, it was identified at the appropriate capture test line (Figure 4B). However, signal intensity drops with more target antigens present as availability of gold nanoparticles for each test band decreases. Thus, mixing the targets in all permutations yielded fainter yet still visible bands as anticipated, demonstrating proof-of-concept multiplexed detection (Figure 4B).

**Figure 4.**
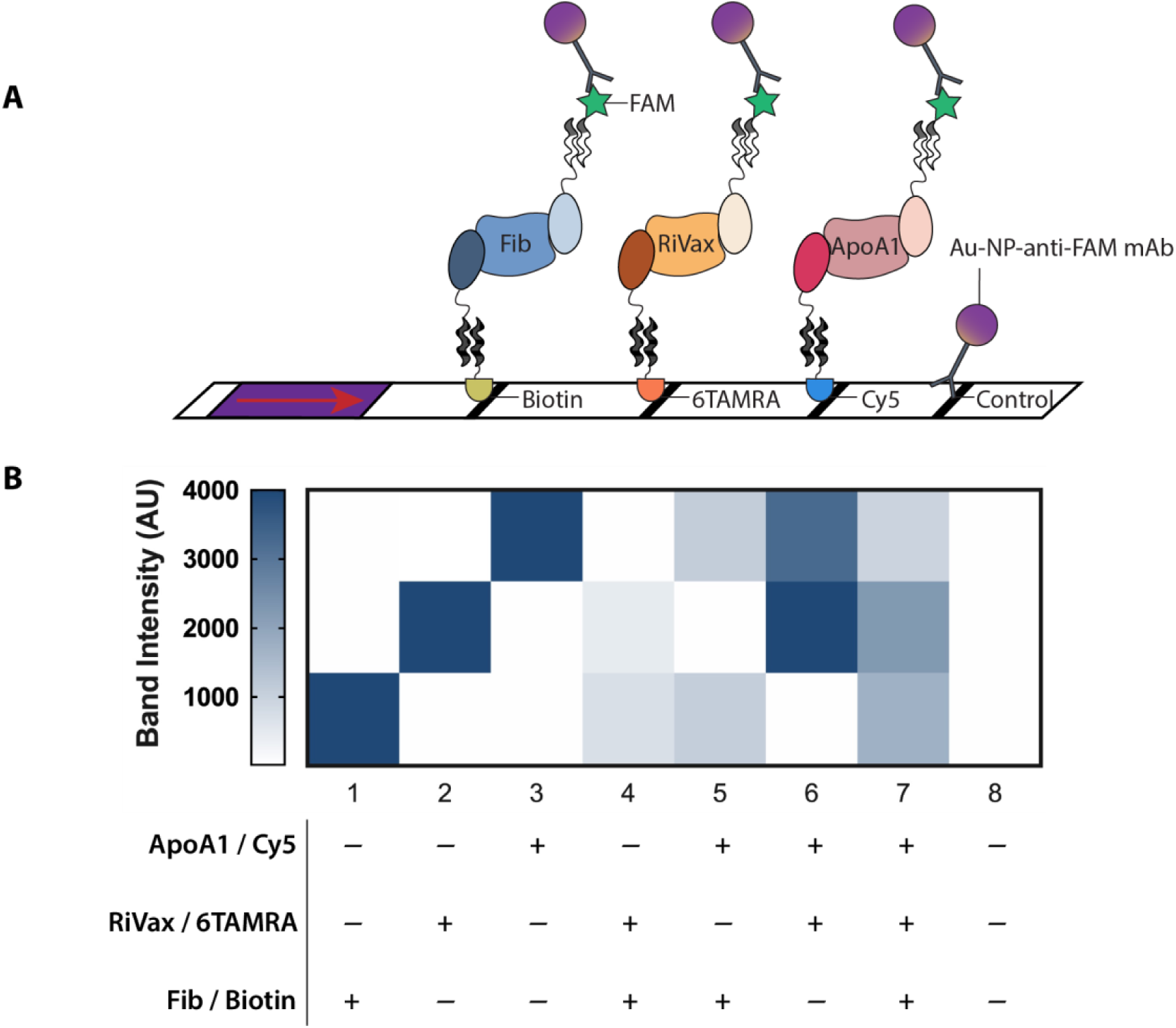
Multiplexed detection of RiVax, ApoA1 and Fibrinogen. (A) Schematic illustration of a multiplexed lateral flow assay strip for all target antigens. (B) Heat map of band intensities for target antigens mixed in various combinations to run on lateral flow strips containing multiple test lines. Results are representative of n=3 independent replicate experiments.

Extending evaluation to a real-world context, we probed RAMONA’s performance in detecting protein biomarkers in human saliva samples. Initially, when untreated and undiluted saliva was used for RAMONA, only 100 nM spiked Fib but not endogenous Fib could be detected using VF1 and VF2 (Supplementary Figure 3). This was not unexpected, as RAMONA’s limit of detection for Fib is near the typical concentration levels of endogenous fibrinogen in saliva.^38^ Nonetheless, this underscored the robustness of RAMONA against pH variations^39,40^, matrix effects^41^ and proteases^42,43^ in saliva. Following 10-fold concentration from saliva using a protein precipitation step, we successfully detected endogenous ApoA1 using Nb11-Nb19, and endogenous Fib using VF1-VF2 sandwich pairs. The performance of RAMONA was benchmarked by comparing against protein detection results from ELISAs. Figures 5B and 5E show pair-wise results for each saliva sample from the two assays detecting ApoA1 and Fib. Pearson correlation coefficients for both targets indicate strong and positive linear correlations (*r* = 0.772 and 0.961 for ApoA1 and Fib, respectively) between ELISA and RAMONA (Figure 5C, 5F). A higher binding affinity of VF1-VF2 for Fib than Nb11-Nb19 for ApoA1, as seen from Figures 3C and 3D, likely contributed to a stronger assay correlation for Fib than ApoA1. While RAMONA sacrifices some sensitivity compared to ELISA, it delivers semi-quantitative results in a fraction of the time – RAMONA can be performed in under 30 minutes with no washing steps, compared to an ELISA which requires 4-6 hours and laborious washing in between incubation steps (Figure 5A).

**Figure 5.**
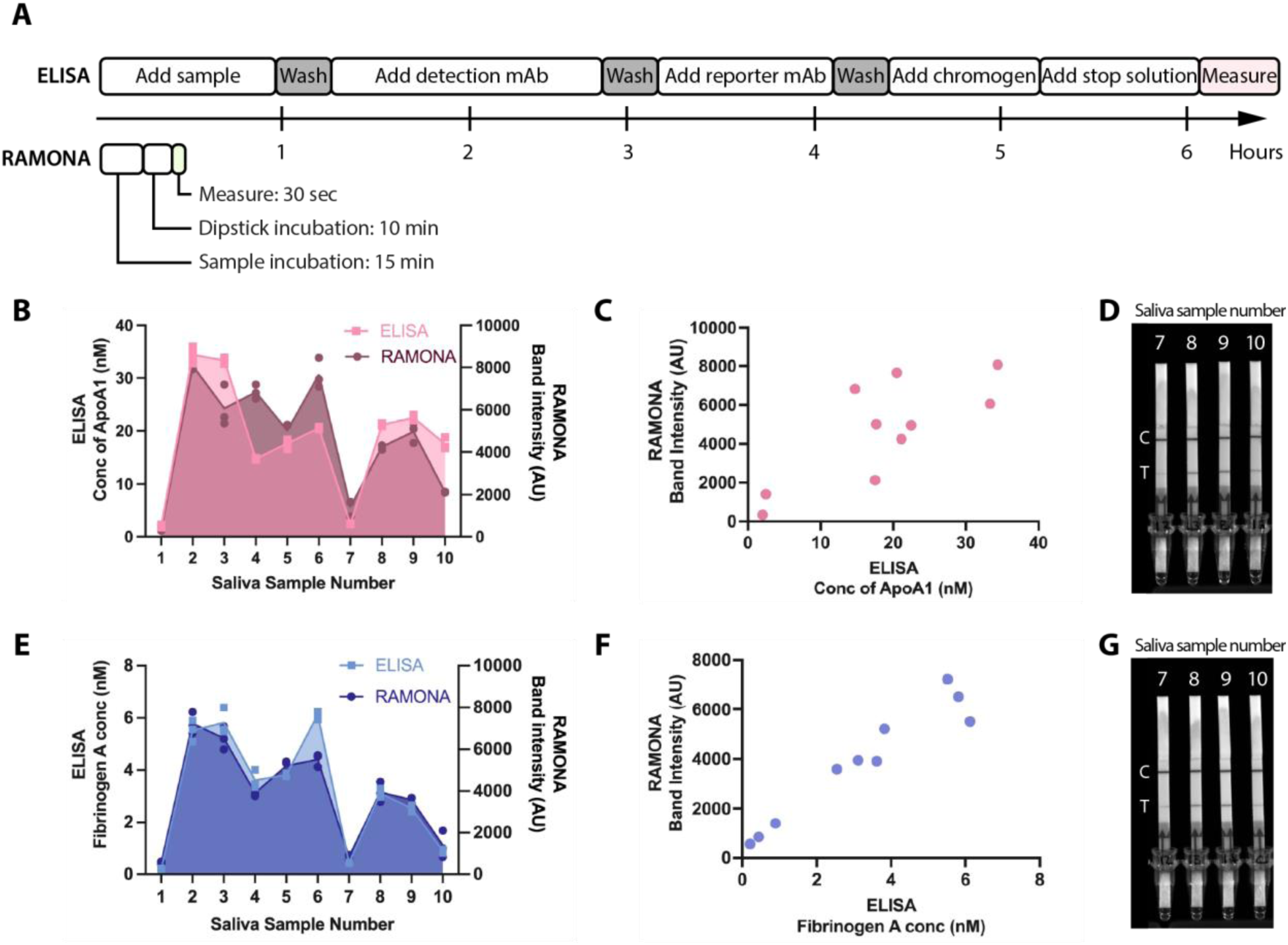
Strong correlation between RAMONA and ELISA detection methods. (A) Schematic comparing the approximate time required for an ELISA versus a RAMONA. (B) Comparison of the concentration of ApoA1 calculated from ELISAs versus the band intensities analyzed from RAMONA LFAs for 10 fresh saliva samples. Untreated saliva was diluted for ELISAs, while 10x concentrated, protein precipitated saliva was used in RAMONAs. Results show n=3 independent replicate experiments. (C) Correlation between mean ELISA and RAMONA results for the same 10 fresh saliva samples in (B), with a Pearson correlation coefficient of 0.772 for ApoA1 (*p*=0.0090). (D) Representative RAMONA LFA images for a subset of fresh saliva samples #7-10, which were chosen to demonstrate variation in band intensities depending on ApoA1 abundance in each sample. “C” indicates control line, and “T” indicates test line. (E) Comparison of the concentration of fibrinogen calculated from ELISAs versus the band intensities analyzed from RAMONA LFAs for 10 fresh saliva samples. Untreated saliva was diluted for ELISAs, while 10x concentrated, protein precipitated saliva was used in RAMONAs. (F) Correlation between mean ELISA and RAMONA results for the same 10 fresh saliva samples in (E), with a Pearson correlation coefficient of 0.961 for Fib (*p*<0.001). (G) Representative RAMONA LFA images for a subset of fresh saliva samples #7-10, which were chosen to demonstrate variation in band intensities depending on Fib abundance in each sample. “C” indicates control line, and “T” indicates test line.

## Discussion

In this work, we describe RAMONA, a nanobody-based platform for portable protein detection that delivers results in under 30 minutes with robustness against environmental factors. It possesses a wide temperature operating window, making it well-suited for field deployment where the ambient environment can vary dramatically with geography. It is functional even in complex biological matrices, such as 80% untreated saliva, where it effectively detects spiked protein targets. However, it does not provide sufficient sensitivity to detect protein targets in the absence of protein concentration and desalt steps. Following sample processing, RAMONA detects endogenous biomarkers present in saliva and provides strong correlations with biomarker concentrations determined from ELISAs. To obviate the need for protein pre-concentration, signal amplification strategies could be integrated into our platform to enhance sensitivity. Other immunoassays incorporating signal amplification demonstrate exceptional sensitivities, suggesting this approach could further extend the utility of RAMONA for detecting low-abundance biomarkers.^44,45,5^

RAMONA is a unique nanobody-based platform that incorporates plug-and-play modularity encompassing both target detection and multiplexing capabilities.^46^ In this study, we attain simultaneous detection of three distinct protein targets by simply substituting nanobodies, eliminating the need for redesigning or re-optimizing the entire system for each new target. While nanobodies conjugated to antigen-specific antibodies have been utilized for multiplexed fluorescence imaging^47^, to the best of our knowledge, multiplexed detection of proteins using nanobodies has only been demonstrated once previously for the duplex detection of hGH isoforms, a system not easily generalizable to other targets.^48^ By contrast, RAMONA offers generalizability and scalability in protein detection.

Furthermore, its integration with cell-free systems further highlights its potential as a transformative platform for rapid and accessible diagnostics. We expand the repertoire of co-binding nanobodies for antigen targets tested via our cell-free screen which revealed known and novel co-binders. Differences between conventional fermentation-based expression and cell-free expression systems, such as ambient oxidation states, may have affected protein folding or disulfide bond formation in different studies, thereby leading to new discoveries of co-binders in this work.^49,50^ For all co-binding nanobody pairs, column chromatography purification after fermentation-based expression led to increased sensitivity of detection as compared to cell-free-based detection. While cell-free reagents without any DNA was found to produce negligible background signal on LFA (data not shown), it is possible that purifying nanobodies from crude cell-free lysates could further improve performance. However, it is currently cost-prohibitive to purify nanobodies from cell-free systems for point-of-need testing. To our knowledge, RAMONA is the first nanobody-based system to demonstrate a viable cell-free workflow for direct protein detection. The isothermal, room-temperature protocol from production to detection streamlines RAMONA’s deployment in low-resource settings. Integration with cell-free systems facilitates rapid prototyping, as evidenced by prior studies that employed these systems for antibody discovery and selection of nanobodies for microscopy.^51,52^ Interoperability with cell-free systems underscores RAMONA’s potential to accelerate the development of diagnostic assays for emerging threats.

In conclusion, RAMONA represents a significant advancement in antigen detection technology, offering a highly adaptable, portable, and efficient platform for rapid diagnostics. Its unique combination of nanobody modularity, cell-free compatibility, and robustness in diverse conditions makes it a promising foundation for developing deployable diagnostics to combat future diseases and threats, enhancing global public health preparedness.

## Materials and Methods

### 1. Nanobody design and plasmid production

Nanobody sequences against antigen targets were obtained from existing literature and fused to one side of a coiled coil peptide pair from a published library via a flexible glycine-serine (GS) linker.^21^ These constructs were then synthesized by Twist Biosciences into a pET32a expression vector (Novagen). Nanobody and coiled coil peptide sequences are available in Supplementary Table 1 and 2, respectively. The following plasmids will be available from Addgene: pET32a-202-P1, pET32a-301-P3S, pET32a-Nb11-P2, pET32a-Nb19-P4S.

### 2. Protein production and purification

Plasmids were transformed into SHuffle® T7 Express Competent *E. coli* cells (New England Biolabs) and streaked on Luria-Bertani (LB, from Sigma Aldrich) agar plates containing 100 µg/mL carbenicillin (Cayman Chemical) for overnight growth at 37°C. To express nanobody-coil fusions, single colonies were picked and grown in 50-100 mL MagicMedia^TM^ *E. coli* Expression media (Thermo Fisher Scientific) using an autoinduction protocol at 30°C for 24 h. Each growth medium was then centrifuged at 4000x*g* for 15 minutes to pellet cells. The supernatant was decanted, and cell pellet frozen at -80°C until ready for protein purification.

All constructs have histidine tags to enable purification by immobilized metal affinity chromatography using HiTrap TALON crude columns (Cytiva). TALON wash and elution buffers were prepared according to the manufacturer’s protocol, with reagents purchased from Sigma Aldrich. Cell pellets were lysed using BugBuster 10X Protein Extraction reagent (Sigma Aldrich) diluted in TALON crude wash buffer. The cell lysates were applied to the TALON crude column on an AKTA pure Fast Protein Liquid Chromatography system (Cytiva) and constructs were eluted using TALON crude elution buffer. The purity and size of selected elution fractions with high A280 values were evaluated using 15% sodium dodecyl sulfate polyacrylamide gel (Bio-Rad Laboratories). Fractions with nanobody-coil fusions were then pooled and desalted using ZebaSpin desalting columns (Thermo Scientific) before their yield was quantified using a Nanodrop. Purified nanobody-coil fusions were stored at -20°C until further use.

### 3. Cell-free production of nanobodies

Nanobody constructs were subcloned into a T7 promoter-driven cell-free optimized backbone (Liberum Biotech, plasmid available with purchase of cell-free lysate) for optimized expression. Polymerase chain reaction was performed using Q5 polymerase (New England Biolabs) on plasmid templates to produce linear amplicons of nanobody constructs (See Supplementary Table 3). Linear DNA for each nanobody was added to individual Juice cell-free lysate mixes (Liberum Biotech) for protein production overnight at room temperature under shaking at 250 rpm. The next day, crude lysates containing nanobodies were functionalized for RAMONA (see below).

### 4. Bioconjugation of synthetic coil peptides

Single-sided coil peptides were made to order with an N- or C-terminal azido group through AbClonal. Depending on the downstream assay type to be interfaced with, various DBCO-containing small molecules (BroadPharm and Integrated DNA Technologies) can be bioconjugated to these coiled coils via copper-free click chemistry (See Supplementary Table 4). Lateral flow is often the most rapid and portable readout of choice, thus a library of FAM- and biotinylated coils were prepared. Briefly, equimolar ratios of an azido-coil peptide and DBCO-PEG-Biotin or DBCO-FAM were incubated overnight at 4°C. Absorption at 310 nm was measured before and after incubation to ensure DBCO depletion, signifying successful click chemistry. These conjugated coils were stored at -20°C until further use.

### 5. RAMONA platform assembly

Nanobody-coil fusions were first functionalized by co-incubation with equimolar amounts of bioconjugated cognate coils at room temperature for 30 minutes. For single antigen detection, protein targets were co-incubated with sandwich nanobodies at various concentrations for 15 minutes at room temperature, unless otherwise stated. For multiplexed detection, equimolar concentrations of nanobodies and target antigens were co-incubated for 15 minutes at room temperature as follows: Fib 600nM, RiVax 300nM, ApoA1 600nM. Concentrations were varied to account for variations in affinities of capture antibodies to their cognate small molecules to yield as even and consistent bands as possible on lateral flow strips. PBS was used to bring total incubation volume up to 15ul if necessary, before lateral flow assays were run. For interface with lateral flow, the reaction mixture was made up to a minimum of 48 µl in volume (usually 1:4 or 1:5 dilution) with ChonBlock Blocking/Sample Dilution ELISA Buffer (Chondrex) before dipsticks (Milenia Biotec) were inserted. A 5- to 10-minute incubation at room temperature followed before the readout was measured using a colorimetric imager. ImageJ software was used to quantify test band intensities.

Human Apolipoprotein A1 was purchased from ACROBiosystems, human Fibrinogen (Fibrinogen3) was purchased from Enzyme Research Labs, and RiVax was a gift from Dr. Alex Miklos and Dr. Dan Phillips from DEVCOM CBC.

### 6. Saliva collection and treatment

Saliva was collected via passive drool into unlabeled 50 ml Falcon tubes (Corning) to de-identify samples. Participants were asked to drink one cup of water 30 minutes prior, and then refrain from eating or drink until after donation. Samples were arbitrarily numbered to enable comparison between RAMONA and ELISA and were aliquoted and frozen at -80°C until further use.

The Native Protein Precipitation kit (Invent Biotechnologies) was used to isolate proteins from saliva samples, with minor optimizations to the manufacturer’s protocol to maximize protein yield. All steps were carried out at 4°C unless otherwise stated. Saliva samples were thawed on ice completely before treatment. Briefly, after thawed saliva was centrifuge-filtered through a column provided at 16,000xg for 10 minutes, the flowthrough was mixed with the precipitating reagent in a ratio of 1:2. This mixture was incubated for an extended period overnight with gentle shaking at 4°C. The next day, samples were centrifuged at 16,000xg for 10 minutes to pellet the precipitated protein. The supernatant was discarded, and the pellet was reconstituted in PBS at a tenth of the initial saliva volume added. Samples were then desalted using ZebaSpin Desalting Columns (Thermo Scientific) at room temperature and stored at -20°C until further use.

### 7. ELISA

Human Apolipoprotein A-I (ApoA1) AssayMax ELISA Kit (AssayPro) and Human Fibrinogen ELISA Kit - high sensitivity (AbCam) were purchased for ELISA tests. Saliva samples were serially diluted and used for testing in triplicate according to the manufacturer’s protocols. Calibration curves were graphed and interpolated in GraphPad Prism to calculate endogenous protein biomarker levels for saliva samples. GraphPad Prism was also used to plot mean values of biomarker levels of RAMONA against ELISA and calculate Pearson r correlation statistics.

## Supporting Information

Additional experimental details, materials and methods (PDF and XLSX).

## Acknowledgements

This work was supported by Defense Advanced Research Projects Agency (DARPA) funding (Contract No. N66001-23-2-4042) and Boston University startup funds to AAG. NRK was supported by the Dial-a-Threat: Antigen program through the Defense Threat Reduction Agency Joint Science and Technology Office (DTRA-JSTO Contract No. CB11133). JJM was supported by NIH Training Program in Quantitative Biology and Physiology (5T32GM008764), NIH Training Program in Synthetic Biology and Biotechnology (1T32GM130546), National Science Foundation Graduate Research Fellowship (2234657). The views, opinions and/or findings expressed are those of the authors and should not be interpreted as representing the official views or policies of the Department of Defense or the U.S. Government. The content is solely the responsibility of the authors and does not necessarily represent the official views of the National Institutes of Health.

## Notes

### Competing Interest Statement

The authors have declared no competing interest.

