## Supplemental Figures for "A Rapid and Modular Nanobody Assay for Plug-and-Play Antigen Detection"

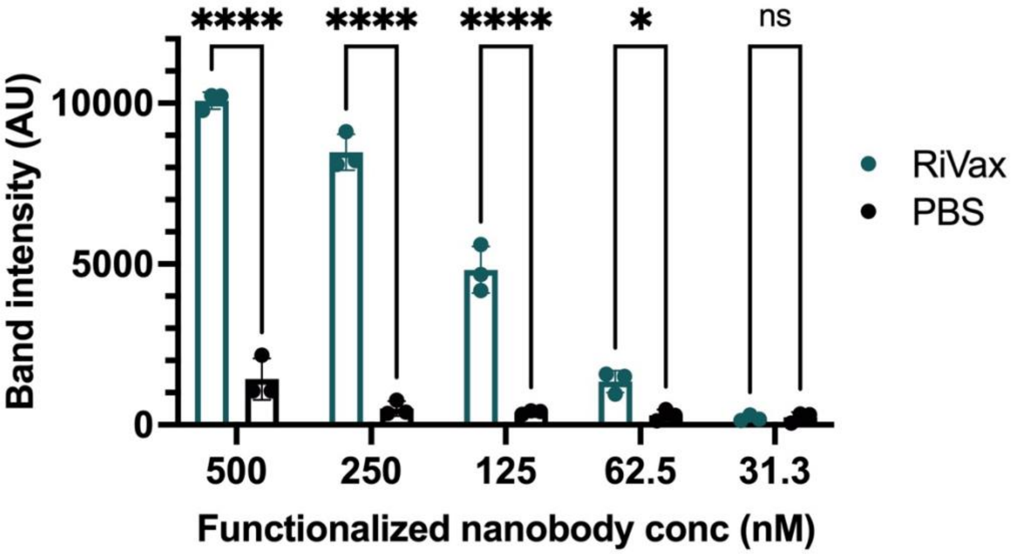


**Supplementary Figure 1.** Titrated concentrations of functionalized RAMONA nanobody-coiled coil conjugates co-incubated with an equimolar ratio of RiVax or PBS for 15 minutes at room temperature before addition to lateral flow strips. Error bars indicate standard deviation across n=3 independent replicate experiments, and statistical significance was determined by two-way ANOVA (adjusted *p* value <0.0001 is denoted by ****, 0.0001 to 0.001 by ***, 0.001 to 0.01 by **, 0.01 to 0.05 by *, and ≥0.05 by ns).

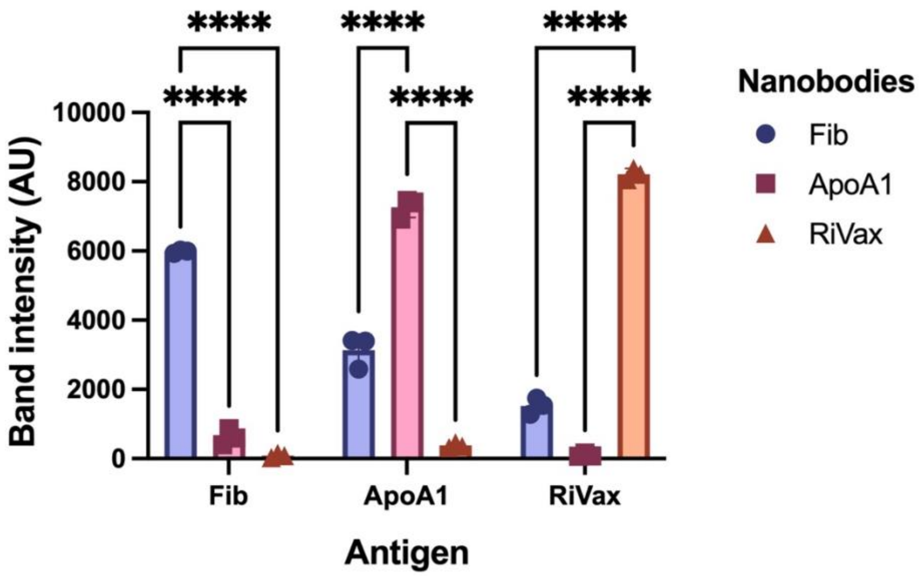


**Supplementary Figure 2.** Test for cross reactivity of nanobodies for other targets. 100nM of functionalized nanobodies were co-incubated with various target antigens in separate reactions at equimolar ratios for 15 minutes at room temperature before addition to lateral flow strips. Error bars indicate standard deviation across n=3 independent replicate experiments, and statistical significance was determined by two-way ANOVA (adjusted *p* value <0.0001 is denoted by ****, 0.0001 to 0.001 by ***, 0.001 to 0.01 by **, 0.01 to 0.05 by *, and ≥0.05 by ns).

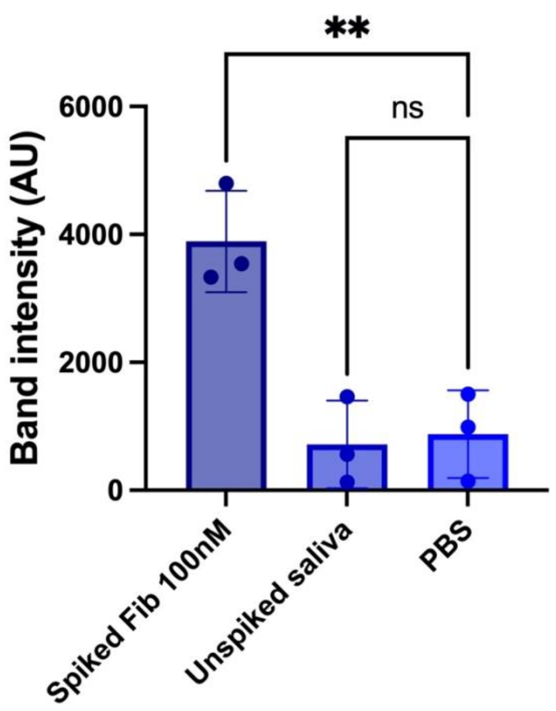


**Supplementary Figure 3.** Compatibility test to determine if 100nM of RAMONA nanobody-coiled coil conjugates VF1 and VF2 could detect spiked or endogenous Fib in untreated and undiluted fresh saliva. Functionalized nanobody-coiled coil conjugates were co-incubated with 83.3% (v/v) spiked or neat saliva for 15 minutes at room temperature before being addition to lateral flow strips. Error bars indicate standard deviation across n=3 independent replicate experiments, and statistical significance was determined by one-way ANOVA (adjusted *p* value <0.0001 is denoted by ****, 0.0001 to 0.001 by ***, 0.001 to 0.01 by **, 0.01 to 0.05 by *, and ≥0.05 by ns).
